# RNAAgeCalc: A multi-tissue transcriptional age calculator

**DOI:** 10.1101/2020.02.14.950188

**Authors:** Xu Ren, Pei Fen Kuan

## Abstract

We introduced RNAAgeCalc, a versatile across-tissue and tissue-specific transcriptional age calculator. We utilized GTEx database to identify 1,616 common age-related genes based on meta-analysis of transcriptional age signature across multi-tissues. Additionally, tissue-specific age-related genes were obtained from differential expression analysis on individual tissues. By performing across-tissue transcriptional age prediction, we showed that our 1,616 common age-related genes outperformed other prior age related gene signatures. Furthermore, we utilized TCGA database to demonstrate that the transcriptional age acceleration computed from our within-tissue predictor was significantly correlated with mutation burden, mortality risk and cancer stage. RNAAgeCalc is available at http://www.ams.sunysb.edu/~pfkuan/softwares.html#RNAAgeCalc.

## Background

Aging is among the most complex phenotype and is a well-known risk factor for a myriad of diseases including cardiovascular, diabetes, arthritis, neurodegeneration and cancer [1]. Increasing evidence has pointed to the interactions between genetics, epigenetics and environmental factors in the aging process [2]. Over the last decade, there has been a growing body of research in identifying genetic and epigenetic biomarkers of aging to decipher the molecular mechanisms underpinning disease susceptibility. For example, the genome-wide association studies (GWAS) have identified genetic loci associated with longevity and several aging-related diseases [3, 4, 5, 6]. As aging is a multifactorial process determined by the dynamic nature of static genetics as well as stochastic epigenetic variation and transcriptomics regulation, both DNA methylation and gene expression have emerged as promising hallmark for understanding the aging process and its associated diseases.

Numerous estimators have been developed to predict human aging from DNA methylation data profiled on the Illumina Infinium HumanMethylation450K BeadChip. Among the most widely used epigenetic age estimator are the DNA methylation age calculator of Horvath [7] and Hannum et al [8], by regressing chronological age on DNA methylation. Specifically, Horvath [7] derived a multi-tissue and cell types DNA methylation age (DNAm age) estimator across the entire lifespan of human. The predictor was constructed using > 8,000 non-cancer samples from 82 DNA methylation datasets, including 51 healthy tissues and cell types based on elastic net [9] model, a penalized regression statistical framework which retained 353 CpGs in the final model. On the other hand, Hannum et al [8] developed a 71-CpGs DNA methylation age calculator based on the whole blood of 656 human samples aged 19 to 101. While the first generation DNA methylation age estimators including Horvath’s clock and Hannum’s clock were developed based on chronological age, the second generation DNA methylation age estimators were obtained by optimizing the prediction error on phenotypic age derived from clinical attributes associated with mortality and morbidity. This includes PhenoAge [10] and GrimAge [11] which aim to improve prediction of aging related outcomes (e.g., time-to-death, time-to-disease for cancer, Alzheimer’s disease and cardiovascular).

In addition to DNA methylation, changes in gene expression have been shown to be associated with aging and aging-related outcomes [12, 13, 14, 15, 16, 17, 18, 19]. Specifically, de Magalhaes et al. [12] identified 56 and 17 genes consistently over- and under-expressed with chronological age, respectively by performing a meta-analysis on 27 microarray datasets from mice, rats and human subjects. Their age specific signature was obtained by first regressing logarithm transformed gene expression on chronological age for each individual microarray dataset. The significance of differential expression in each dataset was determined via a two-tailed F-test, followed by binomial tests to identify genes that were consistently over- or under-expressed across datasets. This study was based on microarray data and 23 out of 27 datasets were from mice or rats subjects, potentially limiting the transferability of the derived age related gene signature to human datasets. As shown in the Results section below, the gene signature of de Magalhaes’s resulted in biased prediction of RNA age based on human RNA-Seq datasets. A closely related work was the development of the GenAge (version 19) database of aging-related genes, including 307 genes potentially related to human aging [13]. Unlike majority of the gene signatures which were typically derived from statistical models (e.g., study specific association analysis or meta-analysis), the genes in GenAge were manually curated by summarizing the biological properties of genes from > 2,000 references on human and animal aging studies across different tissues.

Besides the de Magalhaes et al. [12] signature and GenAge which included across-tissue age-related genes, there were several gene signatures derived for individual tissues. For example, Welle et al. [14] identified 718 probe sites that were related to aging in human muscle using the microarray data from 8 healthy young men and 8 healthy old men. Rodwell et al. [15] identified 985 genes related to aging in human kidney by analyzing the microarray data from 74 healthy kidney with age ranging from 27 to 92 years old, whereas Lu et al. [16] identified 463 aging-related genes in human brain by analyzing the microarray data from 30 samples with age ranging from 26 to 106 years old. Glass et al. [17] identified 1,672 and 188 genes associated with age in skin and adipose tissue, respectively from 856 female twin samples profiled on Illumina Human HT-12 V3 Bead chip.

To the best of our knowledge, the largest meta-analysis to identify age related genes was conducted by Peters et al. [18] from whole blood gene expression of 14,983 human subjects of European ancestry, profiled using microarray platform Illumina Human HT-12 V3 and HT-12 V4 BeadChip. 7,074 samples were used to construct the gene expression signature whereas the remaining samples were used to test the signature. A total of 1,497 genes was significant in both the training and test dataset. The authors further showed that the differences between the predicted transcriptional age and chronological age were associated with biological features related to aging, including blood pressure, cholesterol levels, fasting glucose and body mass index. On the other hand, the most recent transcriptional age predictor was developed by Fleischer et al. [19] based on a novel ensemble linear discriminant analysis (LDA) method using human dermal fibroblast data. Their dataset consisted of gene expression profiled using RNA-Seq from 133 healthy samples with age ranging from 1 to 94 years. The authors further showed that ensemble LDA outperformed other prediction algorithms including elastic net, support vector regression, and linear regression in terms of mean/median absolute difference between predicted age and chronological age. By using leave-one-out cross validation, the authors were able to obtain 4 years median absolute error and 7.8 years mean absolute error in their dermal fibroblast dataset.

Unlike DNA methylation in which several user-friendly software and computer programs are available for predicting epigenetic age across different tissues on the most widely utilized Illumina methylation platform, there were limited transcriptional age predictors and the existing predictors have several pitfalls. First, most of the human transcriptional age predictors were developed based on microarray data and/or limited to only a few tissues. Second, the only predictor constructed using RNA-Seq data was the ensemble LDA predictor [19]. However, this predictor was derived based only on fibroblast data. To date, transcriptional studies on aging using RNA-Seq data across different human tissues was limited. Recognizing the gap in existing research of transcriptional aging based on RNA-Seq data, the aim of this study was twofold, first to identify common age-related genes across tissues; second to construct tissue-specific transcriptional age calculators for understanding how gene expression changed with age in different human tissues. To this end, we utilized a large publicly available RNA-Seq datasets as described in the following sections and developed a transcriptional age predictor for RNA-Seq data. Our transcriptional age predictor is available both as Bioconductor and Python packages RNAAgeCalc, accompanied by a user-friendly interactive Shiny app.

## Results

An overview of the transcriptional age analysis pipeline and comparisons conducted in this study was given in Figure 1.

**Figure 1:**
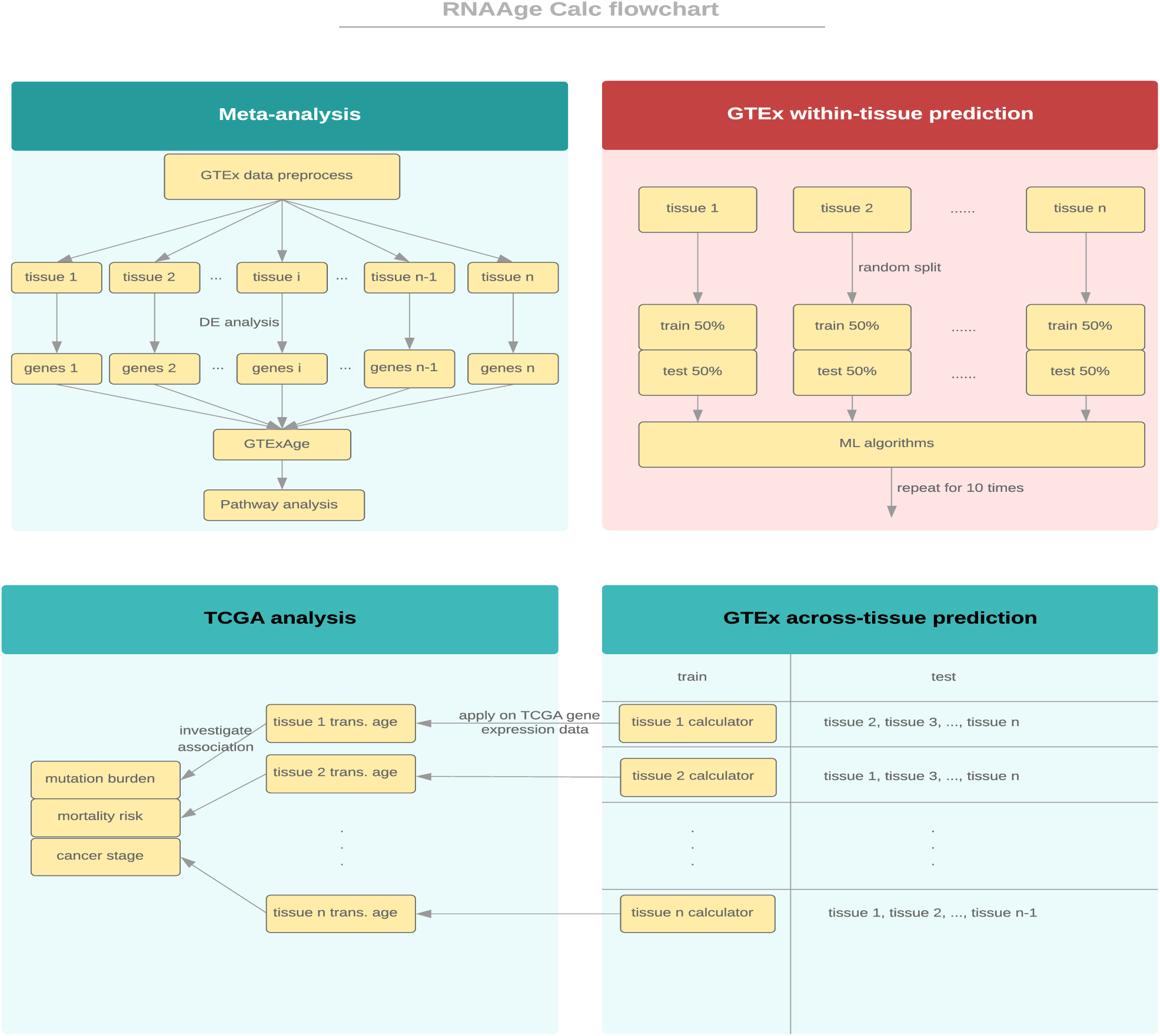
Overview of the transcriptional age analysis pipeline.

### Meta-analysis of age signature across multi-tissues public RNA-Seq dataset

We utilized the RNA-Seq data from the Genotype-Tissue Expression (GTEx) Program [20], a publicly available database to identify across-tissue genes and construct our tissue-specific transcriptional age calculator. We used GTEx V6 release which contained gene expression data at gene, exon, and transcript level of 9,662 samples across 30 different tissues. Since tumor showed notably different gene expression patterns compared to non-tumor [21], the 102 tumor samples from GTEx V6 release were omitted. The remaining samples with complete gender and race information were used in the subsequent analysis. A list of GTEx tissues and summary of non-tumor sample size by gender and race was given in Additional file 1 Table S1. The data preprocessing steps were described in the Methods section.

One pitfall of individual gene expression studies for identifying age-related signatures was the low overlap between the gene lists across different tissues [22, 17]. Meta-analysis of gene expression for aggregating information from different datasets is a useful approach to identify weak genetic signals and has been shown to be powerful in cancer studies [23]. In this paper, we performed a meta-analysis on the different tissues from the GTEx datasets, aiming at identifying common age-related gene expression signatures across the different tissues. After preprocessing, a total of 26 tissues across 9,448 samples were included in our analysis. A summary of number of genes, number of significant genes associated with age under different FDR cut-offs was provided in Additional file 1 Table S2. Certain tissues (e.g. colon, brain) showed strong signal with a high proportion of differentially expressed genes whereas other tissues including small intestine, pancreas, pituitary exhibited relatively weak signal, with only a small proportion of differentially expressed genes. At FDR < 0.05, we obtained 0.02% - 33.08% genes with positive association and 0% - 36.09% genes with negative association across tissues. On average, 13.91% and 13.85% genes were positively and negatively associated with age, respectively. The differentially expressed genes had little overlap across tissues. Among the 26 tissues analyzed, no gene was common across all tissues. Only one gene (EDA2R) was differentially expressed in at least 20 tissues, supporting that age-related signatures were tissue-specific.

To overcome the low overlap across tissues and to identify common age-related genes across tissues, we adapted the binomial test of de Magalhaes et al. [12]. A total of 1,616 common age-related gene across tissues (gender and race adjusted) were identified at FDR < 0.05, as listed in Additional file 2. The details of our approach was described in the Methods section.

### Pathways associated with common age-related genes

The list of genes which exhibited consistent positive association with age were enriched in GO terms related to plasma-membrane adhesion molecules, response to interferon-gamma, GTPase activity, and type I interferon. The enrichment analysis of genes negatively associated with age identified KEGG terms including proteasome, ribosome biogenesis, RNA transport in eukaryotes, citrate cycle, carbon metabolism, pyruvate metabolism, aminoacyl-tRNA biosynthesis as well as GO terms related to mitochondrial function, metabolic process, RNA processing, ribosome biogenesis, and purine metabolic process. Our results were consistent with the findings of Peters et al. [18] which showed that genes involved in RNA metabolism, ribosome biogenesis, purine metabolism, mitochondrial and metabolic pathways were negatively correlated with age. In addition, genes involved in metabolism and mitochondrial protein synthesis were shown to be down regulated in muscle [14]. The study of aging in human brain [16] also indicated that age-related genes in brain played central roles in mitochondrial function. A full list of significant KEGG and GO terms was provided in Additional file 2.

### Within-tissue age prediction and tissue-specific age-related genes

Previous studies showed that age-related signatures were tissue-specific [22]. In this paper, we obtained similar conclusion for the GTEx datasets (see meta-analysis section). To assess whether RNA-Seq data was able to predict chronological age accurately, we first evaluated the age prediction within tissue. We considered a rich class of machine learning prediction models, including elastic net [9], generalized boosted regression models (GBM) [24], random forest [25], support vector regression (SVR) with radial kernel [26], and ensemble LDA [19], as well as numerous candidate feature sets for each algorithm as described in the Methods section. A summary of each candidate feature set and number of genes in each set was provided in Table 1. Our objective here was twofold, first to evaluate which machine learning algorithm performed the best in terms of age prediction, and second to evaluate whether the within-tissue candidate feature sets had better performance compared to across-tissue candidate feature sets.

**Table 1:** Summary of candidate feature sets.

| candidate feature set | number of genes | type |
| --- | --- | --- |
| DESeq | 1,000 | tissue-specific |
| Pearson | 1,000 | tissue-specific |
| Deviance | 2,000-7,000 | tissue-specific |
| Peters [18] | 1,497 | tissue-specific (blood) |
| all | 14,000-19,000 | tissue-specific |
| GTExAge | 1,616 | across-tissue |
| de Magalhaes [12] | 73 | across-tissue |
| GenAge [13] | 307 | across-tissue |

Additional file 3 summarized the average Pearson correlation, Spearman correlation, median error, mean error comparing the predicted age to chronological age for each prediction algorithm, tissue and candidate feature set, across 10 repetitions. Elastic net outperformed other algorithms as illustrated by the highest correlation and lowest mean and median error for almost all tissues and all candidate feature sets. Although ensemble LDA [19] was developed using dermal fibroblast samples, the prediction accuracy on the GTEx fibroblast data was lower than the other algorithms. The prediction accuracy of this model on other tissues was also lower than the other algorithms. These observations indicated that the ensemble LDA [19] predictor may not be generalizable to predict transcriptional age in other tissues or datasets. As elastic net outperformed the other algorithms, we focused on the age prediction using elastic net in the subsequent analysis. Further comparisons of elastic net and ensemble LDA were provided in Additional file 8.

In the elastic net model, the tissue-specific candidate feature sets based on DESeq and all genes outperformed the across-tissue candidate feature sets in most tissues. Another tissue-specific candidate feature set, namely Peters et al. [18] signature developed using microarray blood samples, performed well on the blood samples in GTEx data. On the other hand, our across-tissue signature GTExAge performed best in adrenal gland, breast, lung, small intestine, and stomach but lower performance compared to tissue-specific candidate feature sets in other tissues. Another across-tissue candidate feature set, namely de Magalhaes et al. [12] signature had low prediction accuracy compared to other signatures, which was partially attributed to the fact that it was developed using a large proportion of non-human samples. Taken together, these results suggested that tissue-specific candidate feature sets performed better than across-tissue candidate feature sets in terms of within-tissue age prediction.

### Across-tissue age prediction

To better understand the generalizability of individual tissue-specific transcriptional age predictors to other tissues, we investigated the performance of across-tissue age prediction. Elastic net model was trained on samples from one tissue, and tested on the other tissues. Each gene signature discussed in the Methods section was applied and the Pearson correlation between predicted age and chronological age in test tissue was calculated. The correlation matrix was provided in Additional file 4, where the row represented the training tissue whereas the column represented the test tissue. For each predictor, we calculated the average Pearson correlation across the tissues tested. We then averaged the correlation across all the predictors to evaluate the performance of each candidate feature set. Table 2 showed the average correlation and weighted average correlation by sample size. Our across-tissue feature set GTExAge had the highest correlation, followed by the tissue-specific feature set based on all genes. These results suggested that our across-tissue feature set GTExAge was better than tissue-specific feature sets in across-tissue age prediction.

**Table 2:**
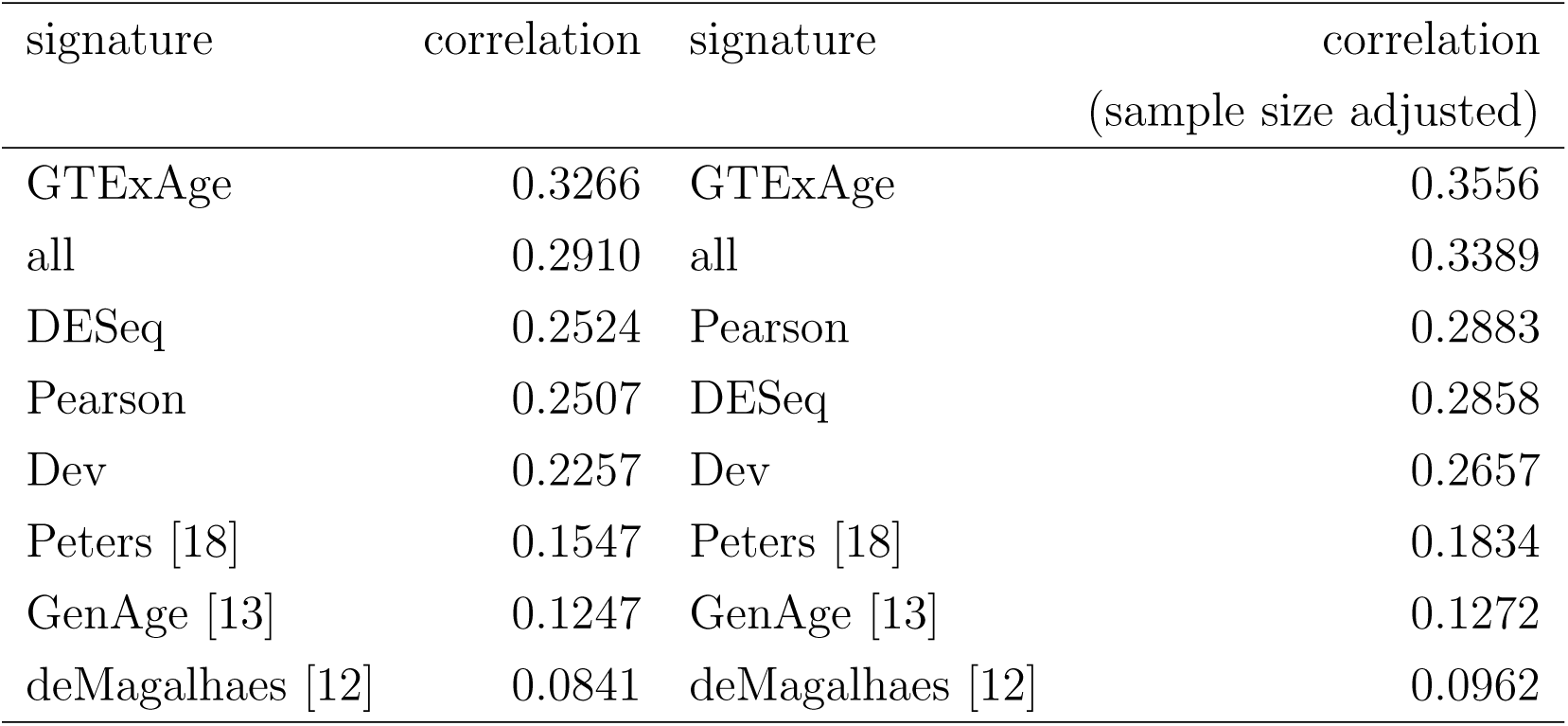
Performance evaluation based on average Pearson correlations across tissues for each candidate feature set.

| signature | correlation | signature | correlation<br>(sample size adjusted) |
| --- | --- | --- | --- |
| GTEXAge | 0.3266 | GTEXAge | 0.3556 |
| all | 0.2910 | all | 0.3389 |
| DESeq | 0.2524 | Pearson | 0.2883 |
| Pearson | 0.2507 | DESeq | 0.2858 |
| Dev | 0.2257 | Dev | 0.2657 |
| Peters [18] | 0.1547 | Peters [18] | 0.1834 |
| GenAge [13] | 0.1247 | GenAge [13] | 0.1272 |
| deMagalhaes [12] | 0.0841 | deMagalhaes [12] | 0.0962 |

The heatmap of Pearson correlation between predicted age and chronological age in each tissue based on GTExAge candidate feature was given in Figure 2. The heatmaps of all genes and DESeq candidate features were given in Additional file 1 Figure S1 and S2. Pairs of tissues showing higher correlation were partially attributable to the tissue lineages and functional similarities [27]. For example, transcriptional age predictor trained on adipose tissue predicted transcriptional age in blood vessel with high correlation (*r*=0.65 in GTExAge signature) due to the similarity in anatomic and function of these two tissues [28].

**Figure 2:**
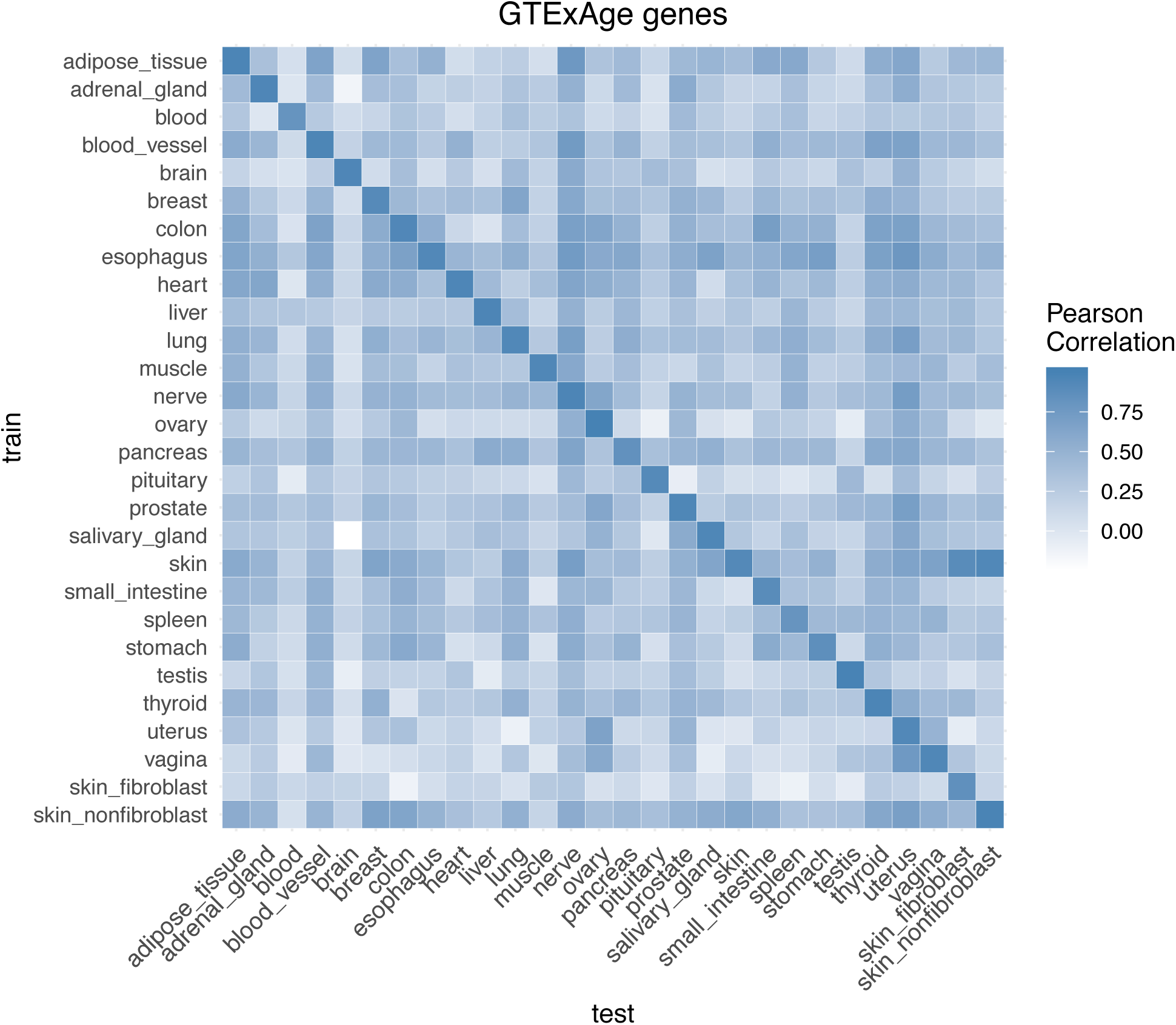
Heat-map of Pearson correlation matrix between predicted age and chronological age (based on GTExAge genes).

### Racial effect on transcriptional age predictor

To evaluate the effect of race on age prediction, we repeated the analysis of the meta-analysis, within-tissue prediction, and across-tissue prediction using only subsets of White samples which made up the racial majority in GTEx database. The predictors were trained on the White samples only and tested on White and non-White samples respectively.

For the within-tissue prediction, the White samples were also divided into 50%-50% training-test set. The predictors were built on the training set and evaluated on the test set as well as on the non-White samples. Additional file 5 summarized average Pearson correlation, Spearman correlation, median error and mean error in each tissue. The predictors tested on White samples had higher correlation and lower error compared to non-White samples in almost all tissues, which indicated the age-related genes were associated with racial differences.

For the across-tissue prediction, elastic net model was constructed using White samples in each tissue. The model was then tested on the White and non-White samples of all the other tissues respectively. The correlation matrix on the White and non-White samples (based on GTExAge candidate feature) was provided in Additional file 6. Figure 3 compared the average correlation on the test sets across all the predictors. The Wilcoxon rank sum test indicated that the difference in the average correlation between the two groups was significant (*p* < 0.05).

**Figure 3:**
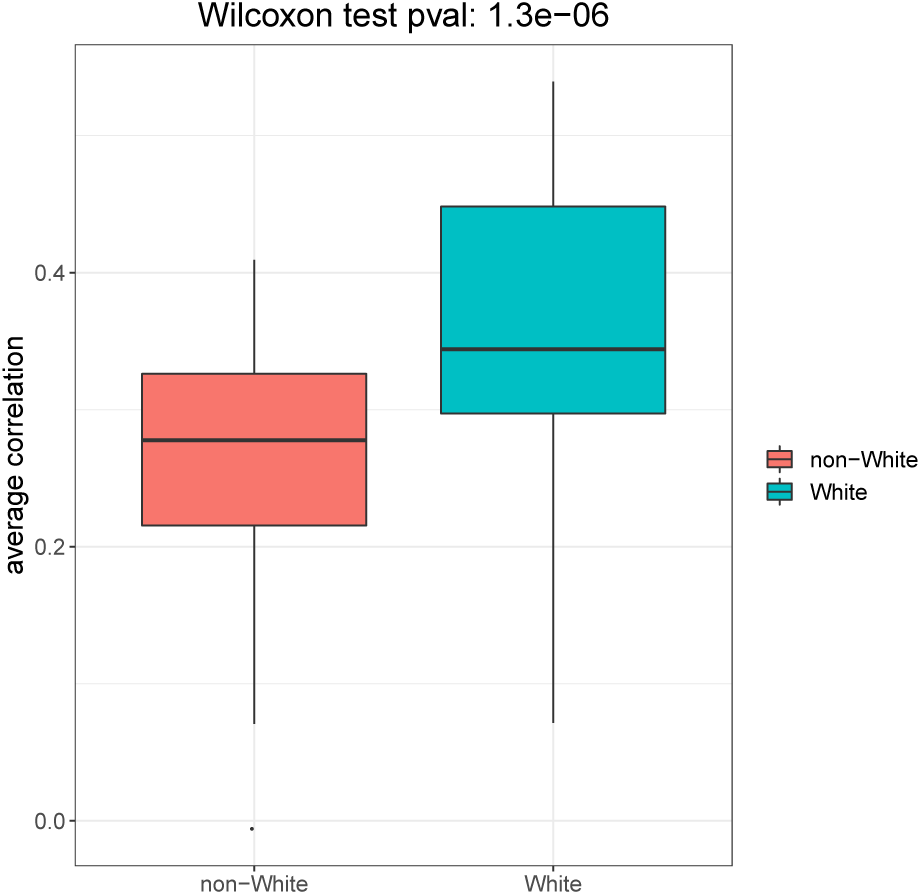
Comparison of the prediction on White and non-White samples (based on GTEx-Age genes).

Both the within-tissue and across-tissue prediction suggested that transcriptional age predictor was racial-dependent. Thus, in our transcriptional age calculator, we provided the option for computing transcriptional age based on models trained on GTEx White samples only.

### Associations with prior aging candidate genes

Here we investigated the intersection between our transcriptional age signature to candidate genes identified from previous studies. First, we compared our tissue-specific signature to prior tissue-specific signatures as summarized in Table 3, Table 4 and Figure 4.

**Table 3:** Overlap between tissue-specific genes in GTEx and prior aging candidate genes.

| prior signature | tissue | original number<br>of genes | number of genes<br>in GTEx | number of genes<br>significant in GTEx* |
| --- | --- | --- | --- | --- |
| Welle et al. [14] | muscle | 479** | 258 | 113 |
| Lu et al. [16] | brain | 463 | 313 | 232 |
| Glass et al. [17] | adipose tissue | 188 | 166 | 100 |
| Glass et al. [17] | skin | 1,672 | 1,334 | 410 |
| Peters et al. [18] | blood | 1,497 | 1,404 | 714 |

\*Genes were considered significant if p-value was less than 0.05.
\*\*These 479 genes corresponded to the 718 probes reported in Welle et al.

**Table 4:** Comparison of the sign of fold change between GTEx and prior aging candidate genes.

| Lu (brain) [16] p = 2.18e-39 |  |  | Glass (adipose tissue) [17] p = 0.0179 |  |  |
| --- | --- | --- | --- | --- | --- |
| GTEx | + | - | GTEx | + | - |
| + | 170 | 12 | + | 49 | 37 |
| - | 31 | 99 | - | 28 | 46 |
| Glass (skin) [17] p = 5.71e-53 |  |  | Peters (whole blood) [18] p = 8.96e-14 |  |  |
| GTEx | + | - | GTEx | + | - |
| + | 446 | 211 | + | 328 | 303 |
| - | 156 | 456 | - | 237 | 502 |

**Figure 4:**
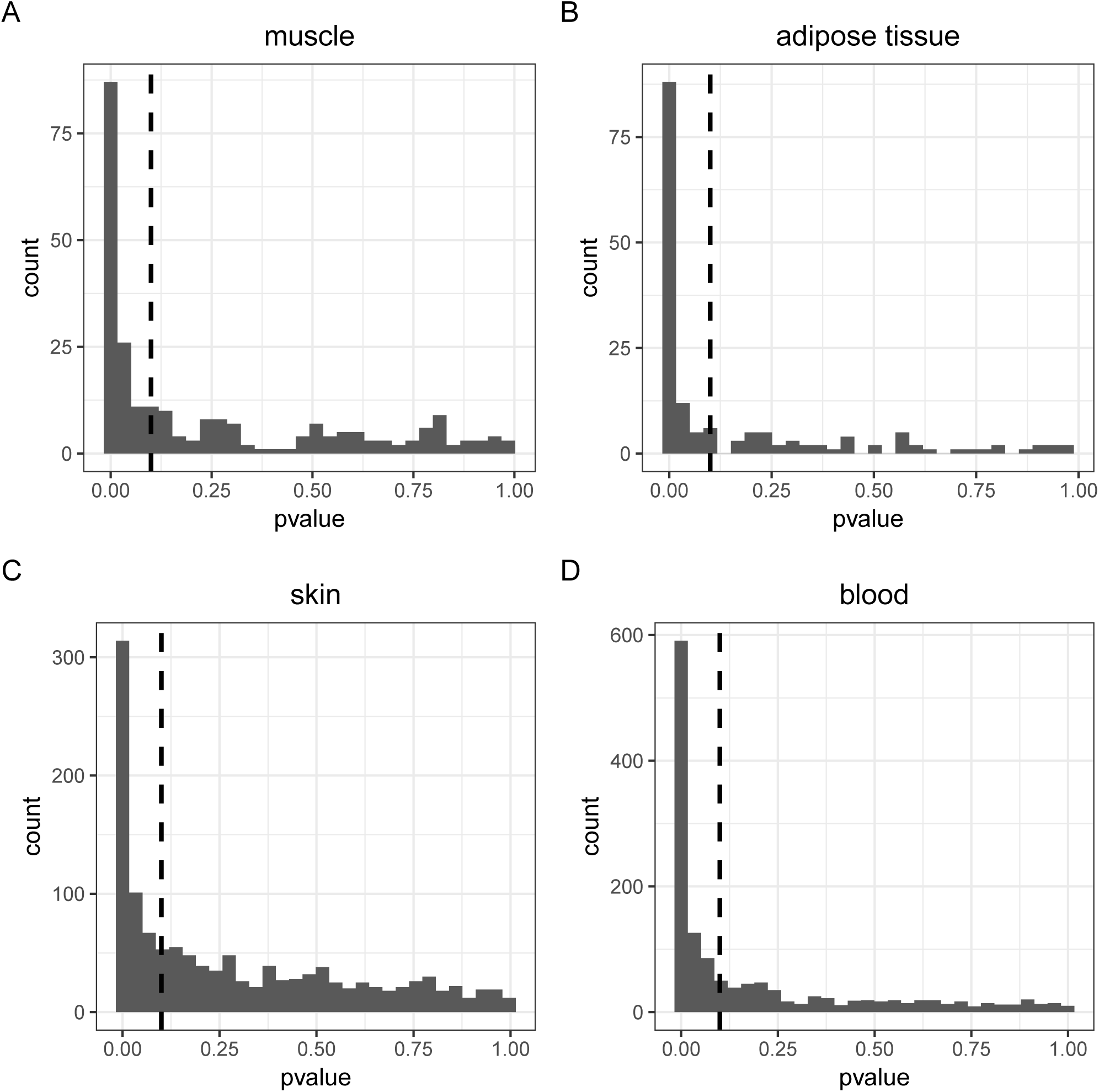
Histogram of p-values of prior candidate genes in GTEx data computed from DESeq2. Dashed line corresponded to p-value 0.1.

In general, our tissue specific signatures obtained from DESeq analyses were consistent with previous tissue-specific candidate age signatures, except for the comparison with Glass et al. [17] skin signature, which could be attributed to the fact that the skin samples were taken from different anatomic regions. Specifically, the skin samples of Glass et al. [17] were from infra-umbilical whereas the skin samples in GTEx were from suprapubic skin, leg and fibroblast. To investigate the performance of transcriptional age prediction based on these prior aging candidate genes, we performed within-tissue transcriptional age prediction using each of these signatures. Table 5 summarized the comparison of prediction accuracy between DESeq/Pearson gene set and the prior candidate genes. For most tissues, the accuracy of our tissue-specific genes selected by DESeq or Pearson correlation were comparable to the accuracy of prior candidate genes.

**Table 5:**
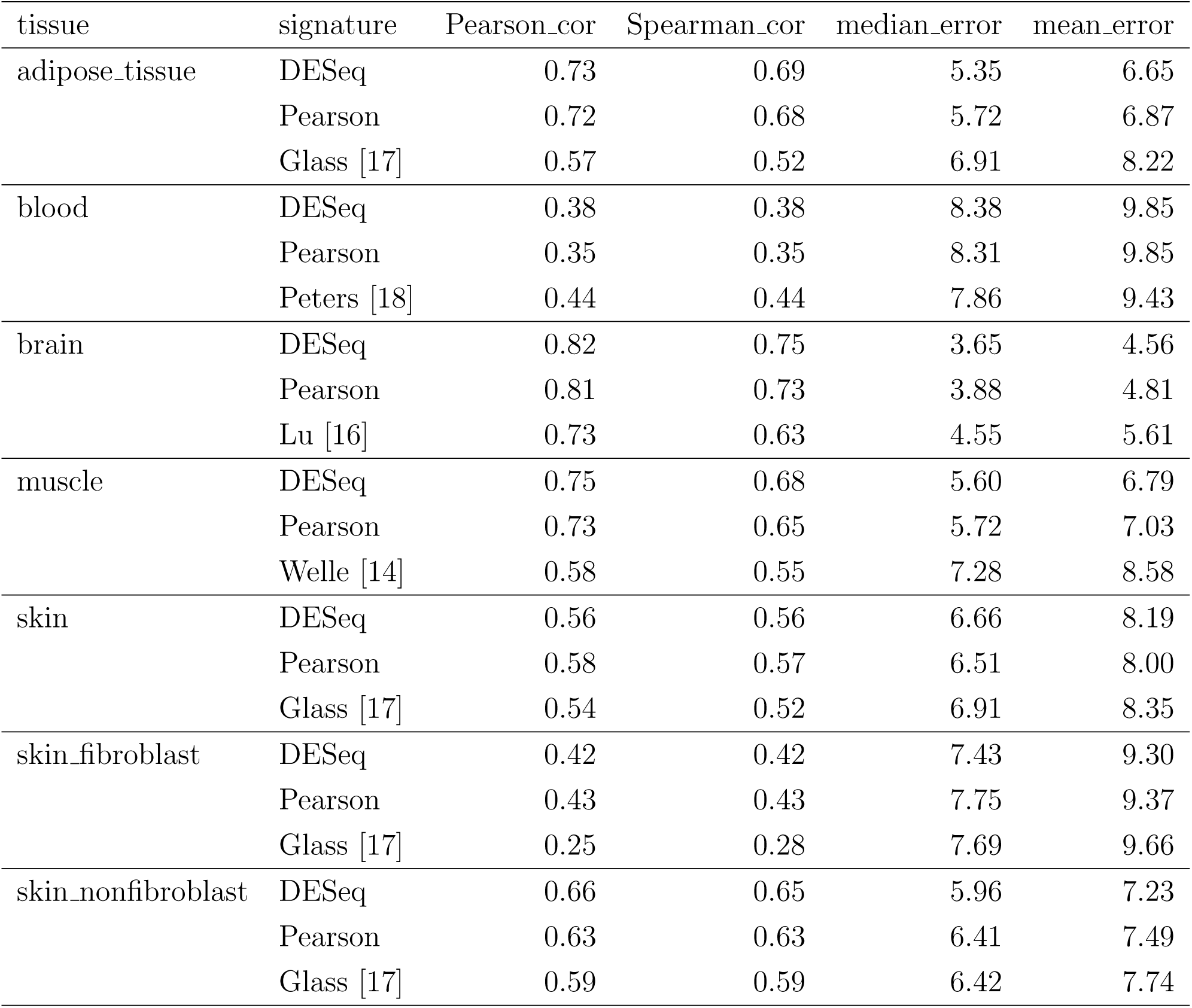
Comparison of GTEx aging genes versus prior candidate genes on within-tissue prediction.

| tissue | signature | Pearson_cor | Spearman_cor | median_error | mean_error |
| --- | --- | --- | --- | --- | --- |
| adipose_tissue | DESeq | 0.73 | 0.69 | 5.35 | 6.65 |
|  | Pearson | 0.72 | 0.68 | 5.72 | 6.87 |
|  | Glass [17] | 0.57 | 0.52 | 6.91 | 8.22 |
| blood | DESeq | 0.38 | 0.38 | 8.38 | 9.85 |
|  | Pearson | 0.35 | 0.35 | 8.31 | 9.85 |
|  | Peters [18] | 0.44 | 0.44 | 7.86 | 9.43 |
| brain | DESeq | 0.82 | 0.75 | 3.65 | 4.56 |
|  | Pearson | 0.81 | 0.73 | 3.88 | 4.81 |
|  | Lu [16] | 0.73 | 0.63 | 4.55 | 5.61 |
| muscle | DESeq | 0.75 | 0.68 | 5.60 | 6.79 |
|  | Pearson | 0.73 | 0.65 | 5.72 | 7.03 |
|  | Welle [14] | 0.58 | 0.55 | 7.28 | 8.58 |
| skin | DESeq | 0.56 | 0.56 | 6.66 | 8.19 |
|  | Pearson | 0.58 | 0.57 | 6.51 | 8.00 |
|  | Glass [17] | 0.54 | 0.52 | 6.91 | 8.35 |
| skin_fibroblast | DESeq | 0.42 | 0.42 | 7.43 | 9.30 |
|  | Pearson | 0.43 | 0.43 | 7.75 | 9.37 |
|  | Glass [17] | 0.25 | 0.28 | 7.69 | 9.66 |
| skin_nonfibroblast | DESeq | 0.66 | 0.65 | 5.96 | 7.23 |
|  | Pearson | 0.63 | 0.63 | 6.41 | 7.49 |
|  | Glass [17] | 0.59 | 0.59 | 6.42 | 7.74 |

Next, we compared our across-tissue signature (GTExAge) to the prior across-tissue signatures, namely de Magalhaes [12], GenAge [13] and Horvath [7] signatures. Horvath [7] signature was based on 353 CpGs from DNA methylation, thus we considered the genes mapping to these 353 CpGs in our comparison. We compared each of these prior across-tissue signature to GTExAge signature by investigating their p-values in the binomial test for identifying common age-related genes across tissues (see Methods section for details). For each gene signature, only a small proportion of genes were significant at *p* < 0.05 (Table 6), indicating that our signature GTExAge provided additional insights into age-related genes.

**Table 6:** Overlap between GTExAge genes and prior aging candidate genes.

| prior signature | original number<br>of genes | number of genes<br>in GTEx | number of significant<br>genes* (positively related) | number of significant<br>genes (negatively related) |
| --- | --- | --- | --- | --- |
| deMagalhaes [12] | 73 | 64 | 12 (18.75%) | 6 (9.4%) |
| GenAge [13] | 307 | 302 | 40 (13.25%) | 42 (13.9%) |
| Horvath [7] | 344** | 267 | 27 (10.11%) | 22 (8.2%) |

\*Genes were considered significant if p-value is less than 0.05 in the binomial test.
\*\*These 344 genes corresponded to the 353 probes reported in Horvath et al.

### Association between transcriptional age and mutation burden in cancer

Since both methylome and gene expression play important roles in aging, to assess whether they complement each other, we compared age-associated methylation to age-associated gene expression in different tissues. We utilized the DNA methylation data and gene expression data of The Cancer Genome Atlas (TCGA) consortium, a rich repository consisting of omics data for multiple types of cancers. The samples with matched DNA methylation data and RNASeq data were analyzed (see Methods section).

The number of mutations per cancer sample (mutation burden) was previously shown to be negatively correlated with DNAm age acceleration [7]. Here, we aimed to determine whether transcriptional age acceleration had significant association with mutation burden. To this end, we calculated the number of somatic mutations for each cancer sample in TCGA dataset. For each cancer type, the Pearson and Spearman correlation between age acceleration residual and logarithmic number of mutations were calculated. Table 7 compared DNAm age acceleration to transcriptional age acceleration, where the transcriptional age was computed using all genes candidate feature set. The results based on DESeq and GTExAge signature as candidate feature set were given in Additional file 1 Table S3. For the transcriptional age acceleration, significant negative associations with mutation burden were observed in adrenal gland (ACC), brain (GBMLGG) and lung (LUAD) cancer (*p* < 0.05). For DNAm age acceleration, negative associations were significant in breast (BRCA), liver (LIHC), lung (LUAD) and prostate (PRAD) cancer. The results indicated that the transcriptional age acceleration and DNAm age acceleration provided complementary information in mutation burden analysis.

**Table 7:** Correlation between age acceleration residual and mutation burden.

|  | transcriptional age acc. (all genes) |  |  |  | DNAm age acc. |  |  |  |
| --- | --- | --- | --- | --- | --- | --- | --- | --- |
|  | Pearson_r | Pearson_pv | Spearman_r | Spearman_pv | Pearson_r | Pearson_pv | Spearman_r | Spearman_pv |
| ACC | -0.2632 | 1.91E-02 | -0.2676 | 1.71E-02 | -0.1194 | 2.95E-01 | -0.0989 | 3.86E-01 |
| BRCA | 0.0135 | 7.27E-01 | 0.0248 | 5.19E-01 | -0.1500 | 1.00E-04 | -0.2322 | 1.29E-09 |
| GBMLGG | -0.2529 | 1.93E-09 | -0.1902 | 7.34E-06 | 0.0842 | 4.88E-02 | 0.0509 | 2.34E-01 |
| COADREAD | -0.0101 | 8.43E-01 | -0.0033 | 9.48E-01 | 0.2352 | 3.47E-06 | 0.0857 | 9.47E-02 |
| ESCA | 0.1891 | 1.06E-02 | 0.0960 | 1.97E-01 | -0.0046 | 9.51E-01 | 0.0008 | 9.92E-01 |
| LIHC | -0.0562 | 2.96E-01 | -0.0480 | 3.72E-01 | -0.1199 | 2.42E-02 | -0.1407 | 8.12E-03 |
| LUAD | -0.2406 | 1.00E-11 | -0.2431 | 6.11E-12 | -0.1039 | 3.82E-03 | -0.0924 | 1.02E-02 |
| OV | -0.6219 | 7.38E-02 | -0.5833 | 1.08E-01 | 0.0785 | 8.41E-01 | -0.0333 | 9.48E-01 |
| PAAD | 0.0240 | 7.62E-01 | 0.0002 | 9.98E-01 | 0.1411 | 7.24E-02 | 0.1194 | 1.29E-01 |
| PRAD | 0.0715 | 1.18E-01 | 0.0751 | 1.00E-01 | -0.1052 | 2.26E-02 | -0.1847 | 5.60E-05 |
| SKCM | 0.1432 | 1.49E-01 | 0.1382 | 1.64E-01 | 0.1645 | 9.69E-02 | 0.1292 | 1.93E-01 |
| STAD | 0.0460 | 3.84E-01 | 0.0588 | 2.66E-01 | 0.1266 | 1.67E-02 | 0.0451 | 3.96E-01 |
| TGCT | 0.0167 | 8.51E-01 | -0.0439 | 6.21E-01 | -0.1883 | 3.26E-02 | -0.1728 | 5.02E-02 |
| THCA | -0.0519 | 2.56E-01 | -0.0239 | 6.00E-01 | 0.0458 | 3.16E-01 | 0.0167 | 7.15E-01 |
| SKCM_metastatic | 0.0631 | 2.32E-01 | 0.0763 | 1.49E-01 | 0.1165 | 2.89E-02 | 0.0953 | 7.42E-02 |
| all_tissues | -0.0124 | 3.76E-01 | -0.0056 | 6.88E-01 | -0.0164 | 2.44E-01 | -0.0630 | 7.39E-06 |

### Association between transcriptional age and mortality in cancer

We evaluated whether transcriptional age was significantly associated with mortality risk in TCGA datasets. For each cancer type, two Cox proportional hazards models were fitted, namely the Cox regression on age acceleration adjusting for chronological age (Mod0a) and Cox regression on age acceleration adjusting for chronological age, stage, gender and race (Mod1a). In Mod1a, gender was not adjusted for breast (BRCA), ovary (OV), prostate (PRAD) and testis (TGCT) cancer. Table 8 showed that transcriptional age was significantly associated with mortality (*p* < 0.05 in Mod1a) in brain (GBMLGG) and pancreas (PAAD) cancer, and marginally associated with mortality (*p* < 0.1 in Mod1a) in skin metastatic samples (SKCM). DNAm age was significantly associated with mortality (*p* < 0.05 in Mod1a) in brain (GBMLGG), stomach (STAD), and skin metastatic samples (SKCM), and marginally associated with mortality (*p* < 0.1 in Mod1a) in esophagus (ESCA) and lung (LUAD). The association between mortality and transcriptional age acceleration showed consistent effect size direction between Mod0a and Mod1a, and vice versa for the association between mortality and DNAm age acceleration. Additional file 1 Table S4 provided the results of transcriptional age constructed using other candidate feature sets.

**Table 8:** Coefficient and p-value of age acceleration residual from Cox regression.

|  | trans. age acc. (all genes) |  | DNAm age acc. |  | trans. age acc. (all genes) |  | DNAm age acc. |  |
| --- | --- | --- | --- | --- | --- | --- | --- | --- |
|  | Coef_Mod0a | PV_Mod0a | Coef_Mod0a | PV_Mod0a | Coef_Mod1a | PV_Mod1a | Coef_Mod1a | PV_Mod1a |
| ACC | -0.0408 | 1.91E-02 | -0.0266 | 5.61E-02 | -0.0181 | 3.34E-01 | -0.0214 | 1.03E-01 |
| BRCA | 0.0044 | 5.27E-01 | -0.0112 | 4.95E-02 | 0.0020 | 7.83E-01 | -0.0070 | 2.22E-01 |
| GBMLGG | -0.0418 | 3.23E-08 | -0.0162 | 1.33E-06 | -0.0418 | 3.23E-08 | -0.0162 | 1.33E-06 |
| COADREAD | 0.0136 | 7.69E-02 | -0.0003 | 9.72E-01 | 0.0026 | 7.68E-01 | 0.0031 | 6.52E-01 |
| ESCA | 0.0049 | 5.37E-01 | -0.0223 | 4.14E-02 | 0.0049 | 5.70E-01 | -0.0214 | 6.91E-02 |
| LIHC | -0.0094 | 9.33E-02 | -0.0008 | 8.84E-01 | -0.0049 | 3.94E-01 | -0.0016 | 7.77E-01 |
| LUAD | -0.0053 | 9.34E-02 | -0.0080 | 3.26E-02 | -0.0032 | 3.14E-01 | -0.0065 | 8.34E-02 |
| OV | -0.0376 | 3.77E-01 | 0.0218 | 7.69E-01 | -0.0376 | 3.77E-01 | 0.0218 | 7.69E-01 |
| PAAD | -0.0228 | 2.03E-02 | -0.0038 | 5.16E-01 | -0.0241 | 1.96E-02 | -0.0012 | 8.47E-01 |
| PRAD | 0.0066 | 9.50E-01 | -0.1944 | 1.24E-01 | 0.0066 | 9.50E-01 | -0.1944 | 1.24E-01 |
| SKCM | -0.0112 | 1.80E-01 | 0.0065 | 6.47E-01 | -0.0115 | 2.21E-01 | 0.0046 | 7.38E-01 |
| STAD | -0.0006 | 8.18E-01 | -0.0116 | 2.42E-02 | 0.0003 | 8.89E-01 | -0.0120 | 1.93E-02 |
| TGCT | 0.0468 | 3.76E-01 | 0.0195 | 7.50E-01 | 0.0513 | 3.77E-01 | 0.0708 | 4.10E-01 |
| THCA | -0.0333 | 2.76E-01 | 0.0050 | 7.80E-01 | -0.0338 | 3.13E-01 | 0.0025 | 8.89E-01 |
| SKCM_metastatic | -0.0063 | 5.23E-02 | -0.0124 | 2.25E-03 | -0.0057 | 8.18E-02 | -0.0127 | 1.94E-03 |

We further evaluated the association between age acceleration and cancer stage. Two linear regression models were fitted for each cancer type, namely regressing age acceleration on stage adjusting for chronological age (Mod0b) and regressing age acceleration on stage adjusting for chronological age, gender and race (Mod1b). Transcriptional age acceleration was significantly associated with stage (Mod1b *p* < 0.05, Table 9 and Additional file 1 Table S5) in adrenal gland (ACC), colon (COADREAD) and liver (LIHC) cancer, whereas DNAm age acceleration was significantly associated with stage in pancreatic (PAAD) and testicular germ cell (TGCT) cancer.

**Table 9:**
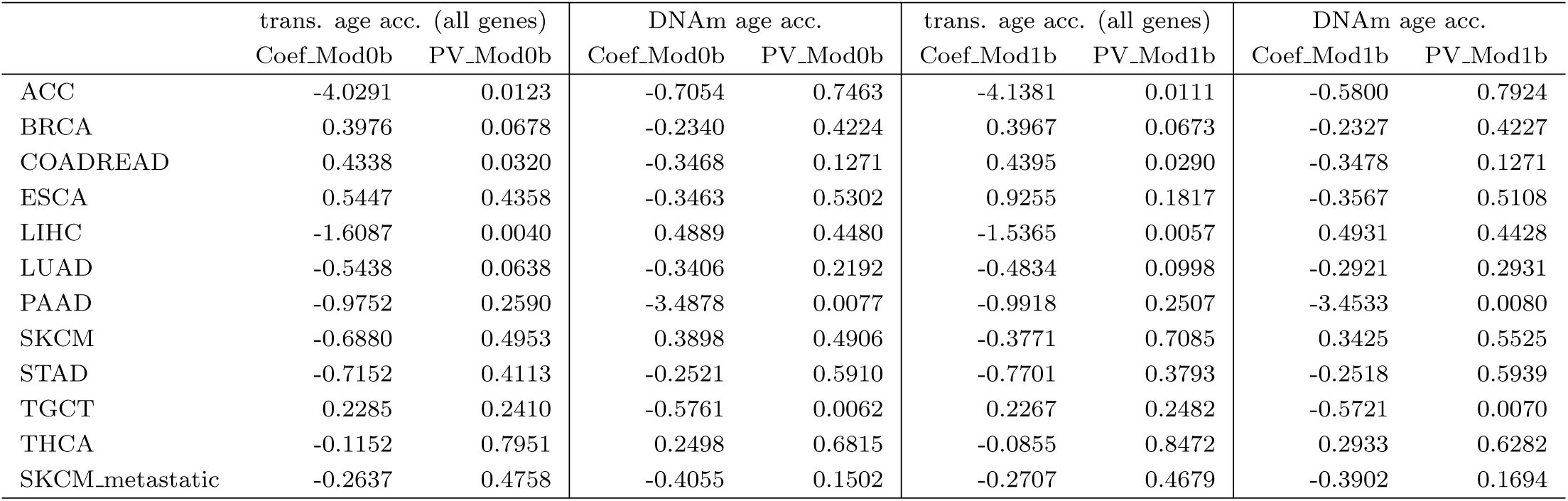
Coefficient and p-value of age acceleration residual versus stage from linear model.

|  | trans. age acc. (all genes) |  | DNAm age acc. |  | trans. age acc. (all genes) |  | DNAm age acc. |  |
| --- | --- | --- | --- | --- | --- | --- | --- | --- |
|  | Coef_Mod0b | PV_Mod0b | Coef_Mod0b | PV_Mod0b | Coef_Mod1b | PV_Mod1b | Coef_Mod1b | PV_Mod1b |
| ACC | -4.0291 | 0.0123 | -0.7054 | 0.7463 | -4.1381 | 0.0111 | -0.5800 | 0.7924 |
| BRCA | 0.3976 | 0.0678 | -0.2340 | 0.4224 | 0.3967 | 0.0673 | -0.2327 | 0.4227 |
| COADREAD | 0.4338 | 0.0320 | -0.3468 | 0.1271 | 0.4395 | 0.0290 | -0.3478 | 0.1271 |
| ESCA | 0.5447 | 0.4358 | -0.3463 | 0.5302 | 0.9255 | 0.1817 | -0.3567 | 0.5108 |
| LIHC | -1.6087 | 0.0040 | 0.4889 | 0.4480 | -1.5365 | 0.0057 | 0.4931 | 0.4428 |
| LUAD | -0.5438 | 0.0638 | -0.3406 | 0.2192 | -0.4834 | 0.0998 | -0.2921 | 0.2931 |
| PAAD | -0.9752 | 0.2590 | -3.4878 | 0.0077 | -0.9918 | 0.2507 | -3.4533 | 0.0080 |
| SKCM | -0.6880 | 0.4953 | 0.3898 | 0.4906 | -0.3771 | 0.7085 | 0.3425 | 0.5525 |
| STAD | -0.7152 | 0.4113 | -0.2521 | 0.5910 | -0.7701 | 0.3793 | -0.2518 | 0.5939 |
| TGCT | 0.2285 | 0.2410 | -0.5761 | 0.0062 | 0.2267 | 0.2482 | -0.5721 | 0.0070 |
| THCA | -0.1152 | 0.7951 | 0.2498 | 0.6815 | -0.0855 | 0.8472 | 0.2933 | 0.6282 |
| SKCM_metastatic | -0.2637 | 0.4758 | -0.4055 | 0.1502 | -0.2707 | 0.4679 | -0.3902 | 0.1694 |

### TCGA matched tumor and normal samples

We applied our tissue-specific predictors on the matched tumor and normal samples from TCGA. Paired t-test and Wilcox test were performed to compare the transcriptional age acceleration residual between tumor and matched normal samples (Figure 5). The tumor samples from breast (BRCA), colon (COADREAD), esophagus (ESCA), prostate (PRAD) and stomach (STAD) cancer showed significant age acceleration (*p* < 0.05) compared to their matched normal samples. We then investigated the aging rate in these paired samples, which was defined as the ratio of transcriptional age to chronological age. As shown in Figure 6, the aging rate was significantly higher in tumor samples compared to matched normal samples in breast (BRCA), colon (COADREAD), esophagus (ESCA), prostate (PRAD) and stomach (STAD) cancer. Since the second generation DNAm age calculator (PhenoAge [10] and GrimAge [11]) were developed to improve prediction of aging related outcomes, we also recomputed the transcriptional age using the genes corresponding to the CpGs of these calculators. The results were provided in Additional file 1 Figure S3 - Figure S8. Overall, DNAm age acceleration based on PhenoAge showed age acceleration in tumor across all cancer types, however the transcriptional age computed based on the genes corresponding to these CpGs showed weaker acceleration pattern, suggesting that the correlation between DNAm and transcription was non-linear and complex [29].

**Figure 5:**
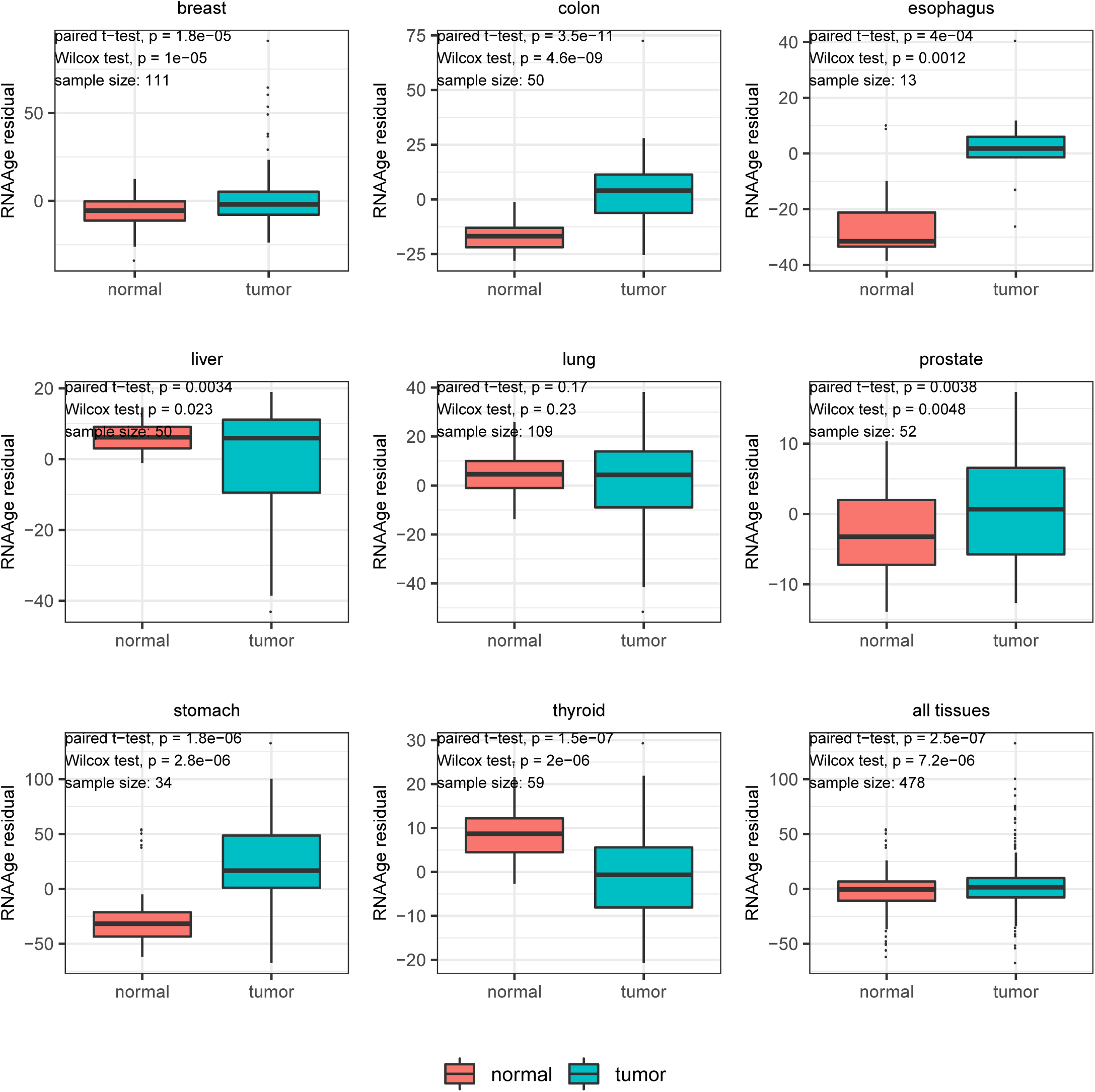
Age predictions on matched tumor and normal samples from TCGA.

**Figure 6:**
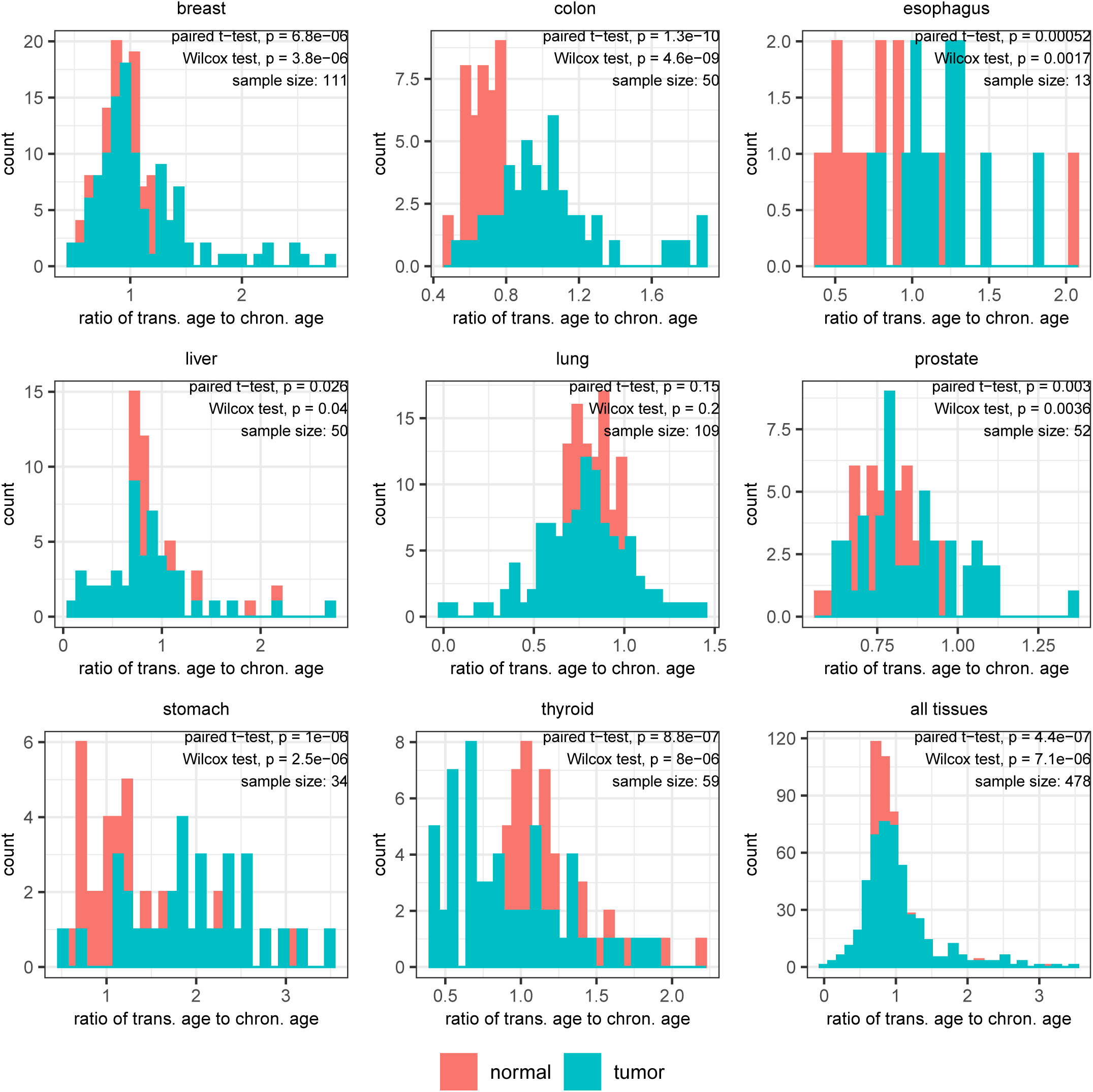
Ratio of transcriptional age to chronological age on matched tumor and normal samples from TCGA.

## Discussion and conclusions

DNA methylation and gene expression were associated with aging and aging-related diseases. A number of calculators to predict DNAm age from human DNAm data profiled on the Illumina Infinium HumanMethylation450K BeadChip have been developed. For gene expression data, although several common age-related genes across tissues as well as tissue-specific signatures have been identified, most of these age-related signatures were developed using either non-human tissues or small sample of tissues. Here, we utilized the gene expression data in the large GTEx database to identify common age-related genes as well as to construct a versatile across-tissue and tissue-specific transcriptional age calculator (RNAAgeCalc). We showed that transcriptional age acceleration was associated with lower mutation burden and lower mortality risk in TCGA cancer samples, and offered complementary information to DNAm age. Our results also indicated that racial difference was associated with transcriptional age. As majority of the samples in GTEx were Whites, future work included extending RNAAgeCalc to non-White samples.

RNAAgeCalc is available both as Bioconductor and Python packages as well as an interactive Shiny app. We anticipate that the calculator will be useful in the development of aging biomarker to advance our understanding on age-related diseases. These insights may ultimately inform development of novel treatments for age-related diseases.

## Methods

### GTEx data processing

To facilitate integrated analysis and direct comparison of multiple datasets, we utilized recount2 [30] version of GTEx data, where all samples were processed using the same analytical pipeline. FPKM values were calculated for each individual sample using `getRPKM()` function in Bioconductor package recount [30]. The benefit of using FPKM instead of raw RNASeq count to build up prediction model was that FPKM had been normalized for the total count and gene length, therefore enabling comparison across different RNA-Seq samples. The recount2 version of GTEx data contained 58,037 genes while the dermal fibroblast data [19] described in the Background section contained 27,142 genes. We studied genes which were measured on both recount2 version of GTEx data and dermal fibroblast data. Genes in recount2 were annotated using Ensembl ID whereas genes in dermal fibroblast data were annotated using RefSeq. We mapped Ensembl ID to RefSeq using Bioconductor package org.Hs.eg.db [31] (version 3.7.0) and only genes with one-to-one map were considered in the analysis, resulting in a total of 24,989 genes.

### Meta-analysis to identify common gene signature

Within each tissue, RNASeq count data was imported from recount2 version of GTEx database and the gene ID subsetting and processing were exactly the same as discussed above. Tissues with fewer than 50 samples were omitted from the analysis as small sample size may lead to conservative and biased results. To avoid the influence of low count genes on the analysis result, genes with more than 20% samples having count per million (CPM) less than one were filtered out. Differential expression analysis were performed on each tissue using DESeq2 [32], adjusting for gender (except for ovary, prostate, testis, uterus, and vagina tissues) and race (White vs non-White). The p-values of differential expression were further adjusted using the false discovery rate (FDR) procedure [33].

We adapted the binomial test of de Magalhaes et al. [12] to identify common age-related genes across tissues. Genes with FDR < 0.05 were considered significantly associated with age. For each individual gene, binomial test was performed with the p-values calculated by the cumulative distribution function:

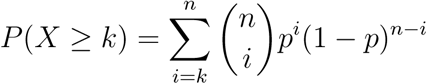

where *n* denotes the total number of tissues, *k* denotes the number of tissues in which the gene was positively (negatively) associated. The parameter *p* was estimated by the average proportion of positively (negatively) associated genes across tissues, resulting in value 13.86% (13.77%). The raw p-values in binomial tests were adjusted using FDR procedure, and statistically significant genes were identified at FDR < 0.05.

### Enrichment analysis of age-related genes

To identify enriched pathways among our 1,616 age-related genes, 297 KEGG pathways [34] and 5,784 Gene Ontologies [35] (minimum and maximum number of genes for each gene set were 15 and 500, respectively) were tested using clusterProfiler software [36]. Hypergeometric tests were performed based on the 828 positively associated genes and 788 negative associated genes, respectively. The p-values from hypergeometric tests were adjusted using FDR procedure and gene sets with FDR < 0.05 were considered significant.

### Transcriptional age prediction via machine learning models

For each individual tissue, we considered several machine learning models including elastic net [9], generalized boosted regression models (GBM) [24], random forest [25], support vector regression (SVR) with radial kernel [26] and ensemble LDA [19]. In all these models, chronological age was the response variable whereas the logarithm transformed FPKM were the predictors. The samples were first randomly split into 50%-50% training-test set, where the prediction algorithms were constructed on the training data and evaluated on the test data. The optimal parameters, namely alpha and lambda in elastic net, number of trees in GBM, cost and gamma in SVR, and bin size in ensemble LDA were selected by 10 fold cross validation in the training set. This 50%-50% training-test split and model evaluation were repeated 10 times. We considered the following candidate gene sets in constructing the prediction models:

i. Differentially expressed genes by DESeq2 [32] (denoted by DESeq). Before training the prediction models, differential expression analysis on age was first performed on the training data. The gene filtering criterion and variables adjusted in differential expression analysis were the same as described in the above section. Instead of using all genes, only the most significant genes from differential expression analysis were used to train the prediction models. Here, we used top *K* differentially expressed genes ranked by the p-values from differential expression analysis. To investigate the influence of the number of genes on prediction accuracy, we considered within-tissue prediction using top 500, 1000, 1,500, 2,000 genes and compared their performances. Additional file 7 showed the prediction accuracy (Pearson/Spearman correlation between predicted transcriptional age and chronological age, median/mean error) versus the number of top significant genes in prediction model. For some tissues the number of top differentially expressed genes did not have a huge impact on prediction accuracy, whereas for other tissues the prediction accuracy increased with the number of top significant genes. To reduce the computation cost, we considered the top 1,000 genes.
ii. Genes highly correlated with chronological age by Pearson correlation (denoted by Pearson). Before training the prediction models, Pearson correlations between the logarithm transformed FPKM and chronological age were calculated on the training set. The correlation coefficients were then sorted by its absolute value in decreasing order, and only the top correlated (either positive or negative) genes were used to train the prediction models. Similar to (i), we also considered top 1,000 correlated genes and a comparison of the number of genes in the model was given in Additional file 7.
iii. Genes have large variance in expression across samples (denoted by Deviance). We adapted the gene selection strategy discussed in [19], in which a gene had at least a *t*_1_-fold difference in expression between any two samples in the training set and at least one sample had expression level > *t*_2_ FPKM to be included in the prediction models. *t*_1_ and *t*_2_ (typically 5 or 10) were the thresholds to control the degree of deviance of the genes. In our analysis, we used *t*_1_ = *t*_2_ = 10 for most tissues. For some tissues with large sample size, in order to maximize the prediction accuracy while maintaining low computation cost, we increased *t*_1_ and *t*_2_ such that the number of genes retained in the model was between 2,000 and 7,000.
iv. The 1,497 age-related genes of [18] (denoted by Peters).
v. All genes after filtering out low count genes.
vi. The 1,616 common age-related genes discussed in the meta-analysis section (denoted by GTExAge).
vii. The 73 common age-related genes of de Magalhaes et al. (denoted by de Magalhaes) [12].
viii. The 307 common age-related genes from the Ageing Gene Database (denoted by GenAge) [13].

The models were evaluated by the Pearson and Spearman correlation between predicted age and chronological age, median absolute error (median error) and mean absolute error (mean error) on the test samples, averaging over the 10 repetitions.

### Associations with prior aging candidate genes

For the tissue-specific signatures, we extracted the genes reported in the original references and took their intersection with the genes available in GTEx. We investigated whether these prior aging candidate genes were significant in our DESeq2 analysis result (Table 3). We then compared the sign of fold change of these prior genes to the sign in our DESeq2 result (Table 4). Fisher exact tests were performed, which showed the signs are highly consistent (*p* < 0.05). The p-values of these prior genes in our DESeq2 analysis result is given in Figure 4.

For each prior across-tissue candidate gene set, we first computed the p-value of each gene using the binomial test (see the meta-analysis section) as a summary measure of evidence the gene held as candidate common age-related gene. We then enumerated the proportion of genes which attained *p* < 0.05 within each prior across-tissue candidate gene set.

### Comparisons of DNAm age versus transcriptional age on TCGA dataset

Illumina Human Methylation 450K annotation data were imported from the Broad GDAC Firehose and the DNAm age were obtained by analyzing the beta value using DNAm age calculator [7, 8, 10]. The TCGA RNASeq data was downloaded and processed from recount2 [30], following the same pipeline as described in the GTEx data processing section. Transcriptional age was obtained by applying the tissue-specific predictors based on all genes, DESeq and GTExAge candidate features on the corresponding tissue in TCGA. For skin cutaneous melanoma (SKCM), the tumor and metastatic samples were analyzed separately. For breast invasive carcinoma (BRCA), only female samples were analyzed. Age acceleration residual was defined as residual from regressing transcriptional age (or DNAm age) on chronological age. The significance of the correlation between age acceleration residual and mutation burden was evaluated by correlation tests. Cox proportional hazards model was fitted on the age acceleration residual and Wald test was performed on the estimated coefficient. Linear regression model was applied to compare age acceleration residual to cancer stage (ordinal covariate) and t-test was performed on the estimated coefficient. Paired sample t-test and Wilcoxon test was used to compare the transcriptional age from matched tumor and normal samples. The p-values < 0.05 were considered statistically significant.

## Acknowledgements

This work was supported in part by the CDC/NIOSH award U01 OH011478. The authors would like to thank Drs. Shannon Ellis and Jeffery Leek for sharing the GTEx metadata.

